# The Queen Gut Refines with Age: Longevity Phenotypes in a Social Insect Model

**DOI:** 10.1101/297507

**Authors:** Kirk E. Anderson, Vincent A. Ricigliano, Brendon M. Mott, Duan C. Copeland, Amy S. Floyd, Patrick Maes

**Affiliations:** USDA-ARS Carl Hayden Bee Research Center, Tucson, AZ 85719; Department of Microbiology, School of Animal & Comparative Biomedical Sciences; University of Arizona, Tucson, AZ, USA 85721; Department of Entomology and Center for Insect Science, University of Arizona, Tucson, AZ, USA 85721

**Keywords:** Honey bees, Acetobacteracaeae, *Bifidobacterium*, *Parasaccharibacter apium*, *Lactobacillus kunkeei*, ileum, aging, core microbiota, bacteria, oxidative stress

## Abstract

**Background:** In social insects, identical genotypes can show extreme lifespan variation providing a unique perspective on age-associated microbial succession. In honey bees, short and long-lived host phenotypes are polarized by a suite of age-associated factors including hormones, nutrition, immune senescence and oxidative stress. Similar to other model organisms, the aging gut microbiota of short-lived (worker) honey bees accrue Proteobacteria and are depleted of *Lactobacillus* and *Bifidobacterium*, consistent with a suite of host senescence markers. In contrast, long-lived (queen) honey bees maintain youthful cellular function without expressing oxidative stress genes, suggesting a very different host environment for age-associated microbial succession.

**Results:** We sequenced the microbiota of 63 honey bee queens exploring two chronological ages and four alimentary tract niches. To control for individual variation we quantified carbonyl accumulation in queen fat body tissue as a proxy for biological aging. We compared our results to the age-specific microbial succession of worker guts. Accounting for queen source variation, two or more bacterial species per niche differed significantly by queen age. Biological aging in queens was correlated with microbiota composition highlighting the relationship of microbiota with oxidative stress. Queens and workers shared many major gut bacterial species, but differ markedly in community structure and age succession. In stark contrast to aging workers, carbonyl accumulation in queens was significantly associated with increased *Lactobacillus* and *Bifidobacterium* and depletion of various Proteobacteria.

**Conclusions:** We present a model system linking changes in gut microbiota to diet and longevity, two of the most confounding variables in human microbiota research. As described for other model systems, metabolic changes associated with diet and host longevity correspond to the changing microbiota. The pattern of age-associated succession in the queen microbiota is largely the reverse of that demonstrated for workers. The guts of short-lived worker phenotypes are progressively dominated by three major Proteobacteria, but these same species were sparse or significantly depleted in long-lived queen phenotypes. More broadly, our results suggest that lifespan evolution formed the context for host-microbial interactions and age-related succession of honey bee microbiota.

## Background

Honey bees (*Apis mellifera*) function as a cooperating group of individuals (colonies) characterized by division of labor [1]. Reproduction is performed by long-lived queen phenotypes while short-lived workers perform a variety of nutrient processing and other tasks that support the reproductive effort [2]. While both longevity phenotypes can result from identical genomes, queens live >10X as long as workers and consume a very different diet [3]. Beginning as newly hatched larvae, queen vs. worker (caste) development is controlled by signaling molecules found in different diets. Pollen exposure halts queen development while royal jelly promotes queen development [4,5]. Nurse workers gorge on pollen to synthesize royal jelly fed to queens throughout their lives. Royal jelly is functionally analogous to mammalian breast milk comprised of a complete diet and antioxidant, antimicrobial and immunoregulatory properties [6,7]. Attributed to caste-specific diets, the phospholipid profile of aging workers becomes increasingly susceptible to oxidative stress, but the queen profile remains stable with age [8]. Consistent with these results, antioxidant gene expression increases in aging workers but not queens [9,10]. Workers live longer when fed the queen diet (royal jelly) as compared to a pollen diet [11]. Collectively, these results suggest that the drastically different lifespans and diets associated with division of labor in honey bees provides a model for mechanisms of diet, aging and microbiota [12,13].

Division of labor in social insects is organized around nutrition and reproduction. In honey bees, this social organization is attributed to the evolutionary repurposing of an egg yolk glyco-lipoprotein (vitellogenin) to serve as nutritional currency throughout the colony [14]. The oldest honey bees leave the hive to forage for nectar, pollen and water. Collected pollen is converted by young workers into two major forms of nutritional currency, one internal; vitellogenin, expressed mostly by abdominal fat body, and one external; royal jelly, shared as social currency among nestmates. In workers, much of the vitellogenin released into the hemolymph is diverted to worker head glands to produce royal jelly [15]. Royal jelly secretions from young (nurse) bees are fed via oral trophallaxis to growing larvae and the queen. In turn, much of the royal jelly fed to queens is converted internally to vitellogenin, to support massive egg production [14]. Vitellogenin is expressed constitutively throughout the queens internal anatomy [9,16]. Like royal jelly, vitellogenin is a multipurpose superfood that functions in immunity, detoxification, oxidative stress, nutrition and longevity [9,16–18]. Older foragers no longer produce jelly, but often beg for and receive small doses from younger nurse bees.

Reproductive division of labor underlies changes in microbiota composition both proximately and ultimately [19,20]. Workers feeding queen vs. worker-destined larvae differ markedly for antimicrobial gene expression associated with royal jelly production in their head glands [21]. Following emergence as winged adults, queen and worker guts are colonized by very different microbiota [20,22,23]. Although highly antimicrobial, the queen’s diet of royal jelly enhances the growth *in vitro* of at least two bacterial species associated with the queen microbiota [7]. Accordingly, the worker phenotype is affected by pollen consumption that occurs concurrent with adult succession of gut microbiota [24]. Experiments with conventionalized honey bee workers and pollen consumption indicate that bacterial fermentation products from recalcitrant pollen shells produced in the gut influence host insulin signaling and the production of vitellogenin [25]. Vitellogenin and life expectancy decrease dramatically as workers transition to foraging and the hindgut microbiota shifts with age [8].

In worker hindguts, fermentation products of gut bacteria are produced according to microbiota structure [25]. A variety of environmental insults can perturb microbiota structure (dysbiosis), altering immune expression, producing oxidative damage and host inflammation [25–28]. Dysbiotic workers suffer developmental deficiencies and early mortality suggesting that the suppression of oxidative stress via microbiota maintenance is critical for gut health and host longevity [25,29]. Similar to gut dysbiosis in response to early life insult, age-associated succession of gut microbiota in worker bees shows increased Proteobacteria with relative decreases in core *Bifidobacterium* and *Lactobacilllus*, the same general results found in many other microbiota models including insects and mammals [12,30]. Unlike workers, the queen does not show greater antioxidant expression with age suggesting that antioxidant function is performed differently or managed by her diet [9]. Vitellogenin hemolymph concentration, constant royal jelly ingestion, and perhaps the microbiota contribute to antioxidant function in long-lived queens.

While research on the worker microbiota has progressed rapidly, little is known of queens. Based on a small sample size and 16S rRNA gene sequencing (amplicons) from whole guts, the queen and worker microbiota differ in taxonomic membership and community structure [19,20,31]. Unlike workers, the early queen gut seems dominated by two distinct species of Acetobacteraceae, *P. apium* and an unnamed species referred to as “Alpha 2.1” [12]. Alpha 2.1 is prevalent in guts of older workers, but *P. apium* occupies a variety of nutrition rich niches associated with honey bees and thrives in the presence of royal jelly [7,32,33]. Capable of gut colonization, *P. apium* is correlated with disease agents in adult bumblebees [34], and in honey bees, implicated in poor worker health, increased mortality, worker gut dysbiosis, and strain dependent effects on larval and pupal survival [12,29,33]. Often occuring with *P. apium*, *Lactobacillus kunkeei* is prevalent/abundant in queens, but like *P. apium*, is also associated with worker disease and dysbiosis [12,35]. *L. kunkeei* is also considered a hive (not a gut) bacterium due to its association with fructose rich niches like honey and honey rich pollen storage [32,36]. Thus, investigations of honey bee microbiota require a careful consideration of social and functional context including host longevity, caste specificity, developmental stage, potential refugia and transmission from nutrition related niches [22,23,37].

Here we test the hypothesis that lifespan differences in a social insect model are associated with age-based microbial succession. We sample known age queens from different backgrounds, and compare our findings to the extensive preexisting characterization of known age workers. We define the aging queen microbiota by deep sequencing four alimentary tract niches that differ in many ways including physiological function, pH and oxygen exposure. To accompany each amplicon library, we determine the absolute numbers of bacteria with qPCR. Finally, we quantify protein oxidation in the fat body tissue of each queen to test the hypothesis that biological age differs from chronological age, and that the accrual of oxidation products in aging queens is associated with species-specific differences in the microbiota.

## Methods

### Queen sampling

Our sampling design distinguished environmental exposure from chronological age. We sampled four different sets of queens; young queens (1^st^ year, n = 31), aged 4-6 months and old queens (2^nd^ year, n = 32) aged 16-18 months. To control for source variation we sampled old and young queens from similar and different backgrounds. The primary model contains two main effects and an interaction effect, asking whether the variation in queen microbiota depends on age, background, or an interaction of both factors.

We sampled a total of 63 queens. Half (n= 32) of these queens were sourced from a large migratory beekeeping operation based in southern California. Referred to as the “CA” source, these Italian queens (*Apis mellifera ligustica*) were purchased from the same queen breeder in different years (mid-March of 2015 and 2016). Both sets of queens were sampled in mid July 2016. Thus “older” CA queens (n = 16) were sampled from colonies that had survived 16.5 months and experienced almond pollination and two seasons of alfalfa pollination in the Imperial valley of southern California. Following almond pollination in 2016,”young” CA queens (n = 16) were introduced via colony splits in March, experienced one season of alfalfa pollination, and were sampled in at 4.5 months of age.

The other half of our sampled queens, referred to as “AZ” queens (n = 31) were sourced from two very different environmental and genetic backgrounds. We sampled “young” AZ queens (n = 15) from the Carl Hayden Bee Research Center in Tucson Arizona. Delivered and installed with 3000 young worker bees (package bees) in early May 2016, these Italian queens (*Apis mellifera ligustica*) were exposed to varied pollen and nectar sources typical of the Sonoran desert, but not intensive agriculture or bulk transportation events. Young AZ queens were sampled in early October, 2016 at 5.7 months of age. In contrast, old AZ queens originated from a Northern migratory beekeeping operation that raises *Apis mellifera carnica* queens. These queens were introduced in April of 2015 to colony splits in the foothills east of Turlock CA following almond pollination. Colonies then experienced the summer in North Dakota making honey, pollinating oilseed crops, sunflowers and canola. Colonies then overwintered in a temperature controlled warehouse in Idaho (November-January), and pollinated almonds in central California (February). The colonies were then delivered to Tucson, Arizona in March of 2016 where they flourished for seven months before queens were sampled in early October at 18 months of age.

All 63 queens were collected into sterile 2.0ml tubes and immediately frozen on dry ice and stored at −20°C for DNA extraction. Queens were dissected under sterile conditions. Four tissue types were extracted to typify the queen microbiota: mouth parts, midgut, ileum and rectum. Mouthparts were unfolded out of the head capsule and detached proximal to the labrum with sterile scissors. Individuals were then pinned through the thorax and the digestive tract was accessed by removing the dorsal abdominal sclerites. The entire digestive tract was removed and floated in 70% EtOH to wash and separate the midgut, ileum, and rectum. The abdominal fat body and attached dorsal sclerites were retained as a single unit to quantify biological age.

### Queen aging assay

As a proxy for biological age, we quantified molecular by-products that cannot be excreted, but accumulate with age in abdominal fat body tissues. In honey bees the accumulation of oxidized proteins (carbonyl groups) in the fat body is recognized as a marker of chronological age [38]. Carbonyl content of total fat body protein homogenates was determined using a commercially available kit (MAK094; Sigma-Aldrich). Briefly, whole fat bodies were homogenized in 600ul of 1X TE buffer. The supernatant was treated with a final concentration of 10 mg/ml streptozotocin to precipitate nucleic acids. The supernatant was decanted then reacted with 2,4-dinitrophenylhydrazine (DNPH) to form stable dinitrophenyl hydrozone adducts. Derivatized proteins were precipitated with trichloroacetic acid and were washed three times with acetone. The samples were resuspended in 100ul of 6M guanidine (pH 2.3). Protein oxidation, expressed as nanomoles of carbonyl groups per milligram of protein was calculated by absorbance at 345 nm relative to the millimolar extinction coefficient of aliphatic hydrozones (22.0 mM^−1^ cm^−1^). The protein content of each sample was determined using a bicinchoninic acid (BCA) assay [39].

### DNA extraction and qPCR

Dissected tissues were placed immediately into 2-ml bead-beating tubes containing 0.2 g of 0.1-mm silica beads and 300 μl of 1X TE buffer. Samples were bead beaten for a total of 2 minutes at 30s intervals. To each sample, 100 μl lysis buffer (20 mM Tris-HCl, 2 mM EDTA, 5 % Triton X-100, 80 mg/ml lysozyme, pH 8.0) was added and the samples were incubated at 37°C for 30 min. DNA was then purified using a GeneJet Genomic DNA Purification Kit according to the manufactures instructions for gram-positive bacteria.

We quantified total bacterial abundance for each of the four tissue types with a real-time PCR (qPCR) assay of 16S rRNA gene copies [40]. A standard curve was generated using a serial dilution of a plasmid standard containing a full length *Escherichia coli* 16S rRNA gene. The assay was validated for use on honey bee-associated bacteria by confirming amplification against individual plasmid templates harboring full length 16S genes corresponding to major gut phylotypes. The qPCR results were expressed as the total number of 16S rRNA gene copies per DNA extraction (200ul volume elution).

### Amplicon pyrosequencing

The V6–V8 variable region of the 16S rRNA gene was amplified using PCR primers 799F (acCMGGATTAGATACCCKG + barcode) and bac1193R (CRTCCMCACCTTCCTC). Amplification was performed using the HotStarTaq Plus Master Mix Kit (Qiagen, USA) under the following conditions: 94 °C for 3 min, followed by 28 cycles of 94 °C for 30 s, 53 °C for 40 s and 72 °C for 1 min, with a final elongation step at 72 °C for 5 min. PCR products were confirmed using a 2% agarose gel. PCR products were used to prepare DNA libraries following Illumina TruSeq DNA library preparation protocol. Sequencing was performed on a MiSeq at the University of Arizona Genetics Core.

### Pyrotagged sequence analysis

Amplicon sequences were processed using MOTHUR v.1.35.0 [41]. Forward and reverse reads were joined using the make.contigs command. After the reads were joined the first and last five nucleotides were removed using the SED command in UNIX. Using the screen.seqs command sequences were screened to remove ambiguous bases. Unique sequences were generated using the unique.seqs command. A count file containing group information was generated using the count.seqs command. Sequences were aligned to Silva SSUREF database (v102) using the align.seqs command. Sequences not overlapping in the same region and columns not containing data were removed using the filter.seqs command. Sequences were preclustered using the pre.culster command. Chimeras were removed using UCHIME [42] and any sequences that were not of known bacterial origin were removed using the remove.seqs command. All remaining sequences were classified using the classify.seqs command. All unique sequences with one or two members (single/doubletons) were removed using the AWK command in UNIX. A distance matrix was constructed for the aligned sequences using the dist.seqs command. Sequences were classified at the unique level with the RDP Naive Bayesian Classifier [43] using a manually constructed training set containing sequences sourced from the greengenes 16S rRNA database (version gg_13_5_99 accessed May 2013), the RDP version 9 training set, and all full length honeybee-associated gut microbiota on NCBI (accessed July 2013). OTUs were generated using the cluster command. Representative sequences for each OTU were generated using the get.oturep command. To further confirm taxonomy, resulting representative sequences were subject to a BLAST query using the NCBI nucleotide database. Diversity indices were generated using the rarefaction.single and summary.single (alpha diversity) commands.

### Statistical analysis

To examine the effect of community size we multiplied the proportional abundance of OTUs returned by amplicon pyrosequencing by the total bacterial 16S rRNA gene copies determined with qPCR for each individual queen and niche. All core bacterial genomes contain four 16S rRNA gene copies except *L. kunkeei* (5), *Bifidobacterium* (2) and *P. apium* (1). Acetobacteraceae Alpha 2.1 (copy number unknown) was designated a value of one, consistent with the copy number of its closest relative, *P. apium* [44]. OTUs representing non-core diversity were summed (Σ OTUs 10-500), corrected for community size and mean 16S rRNA gene copy number (4.2) [45], and used to assess the change in diversity abundance with chronological age and cellular damage. In this case, absolute abundance is extrapolated from relative abundance of amplicons, so remains compositional.

To allow the use of parametric multivariate analyses [46], we converted bacterial abundances to ratios among all OTUs [47] using the software CoDaPack’s centered log-ratio (CLR) transformation [48]. We compared microbial community structure by chronological age and source using a two-way factorial MANOVA and a post-hoc test to compare specific bacteria across conditions. We compared absolute abundance of each bacterial taxon by age without reference to source variation using the Wilcoxon rank sum test. Finally we examined the relationship between carbonyl accumulation and the microbiota in various ways: 1) Using DistLM we test whether the microbiota from each of four distinct tissues is significantly associated with carbonyl accumulation in queens, 2) We examine carbonyl accumulation as a covariate in three separate MANCOVA models, a two-way examining source and age, a one-way examining source, and a one-way examining age, 3) We calculate independent Pearson’s correlations between species-specific CLR scores and log transformed carbonyl data, and 4) We perform principle component analysis, plotting the relationship of bacterial community composition and age associated succession relative to carbonyl accumulation by niche. For the extended queen data set we calculated correlations among the top 200 OTUs using Sparse Correlations for Compositional data algorithm [SparCC:[49]] as implemented in mothur [41]. SparCC is robust for compositional data sets with a low effective number of species [50]. Analyses were conducted in JMP_ v11 (JMP_ 1989–2007) and/or SAS_ v9.4 [51].

We compare our queen results to worker data from a recently published manuscript [19]. As one of three studies sequencing both nurses and foragers, the Kwong *et al.* study is the largest and most robust, and provides whole gut microbiota based on 16S rRNA gene sequences from worker *Apis mellifera; n =* 84 workers; 19 foragers (old) and 65 in-hive bees (young). From this data set we designated eight core gut bacteria representing 95% of OTU abundance based on known samples of whole gut communities in the literature. The remaining 5% OTU abundance from [19] was comprised primarily (83%) of Proteobacteria, occurred with sporadic abundance and prevalence across many worker studies, and was combined as an “OTHER” category to represent low abundance bacteria or signs of dysbiosis. As stated above for queens, we CLR transformed relative abundance measures of workers, and performed a one-way MANOVA on age to compare forager vs. in-hive bee gut microbiotas, calculating post-hoc differences between specific bacterial groups. To compare the queen and worker results we transform our tissue specific queen data to reflect relative abundance values predicted for the whole gut. Tissue specific bacterial cell counts were used to normalize the relative occurrence of bacterial species by queen tissue, then additively produce a single value that represents the expected result of sequencing whole queen guts. These whole gut values highlight differences in abundance and prevalence between workers and queens.

## Results

### Next generation sequencing and qPCR

Next generation sequencing returned 7.2 million quality trimmed reads (400 bp assembled) across the 252 libraries (63 queens X 4 niches). Read coverage was sufficient for all downstream characterization and statistics (Table S1). The queen rectum was represented by 2.4 million reads averaging 38K per library, the ileum by 1.9 million reads averaging 30K per library, the midgut by 1.6 million reads averaging 25K per library, and the mouth by 1.3 million reads averaging 21K per library. The nine most common OTUs (97%) accounted for 98.8% of all reads across the combined niches. Given the low effective number of OTUs, unique OTUs were manually assessed to verify 97% species clusters. Subtracting the rare biosphere (1.2%), these nine OTUs are what we present in figures and use in statistical analyses. All recovered species clusters correspond to previously sampled phylotypes from worker guts or hive materials. Summed across the four niches, the nine most abundant OTUs according to raw amplicon read totals were *L. firm5* (51.3%), *P. apium* (27.1%)*, L. kunkeei* (7.6%), *L. firm4* (6.8%)*, Alpha 2.1* (2.0%), and *Bifidobacterium* (1.5%), S. *alvi* (1.8%) and *G. apicola* (0.3%) and *Delftia* spp. (0.2%). In honey bees, *Delftia* is an unrecognized (novel) species of Burkholdariales that may prove functionally important to host physiology.

Similar to the abundance pattern in worker guts, the queen rectum harbors an average of 121.2M 16S rRNA gene copies per queen, a magnitude more than the ileum (17.9M) or midgut (14.2M). The mouth (1.4 M) contains the least bacteria. Total amplicon reads returned for the mouth, midgut and ileum were significantly correlated with bacterial abundance as determined by qPCR (Table S2). Species dominance in the queen increases with community size in the mouth, midgut and ileum (Table S2). Extrapolating qPCR results to estimate absolute abundance, *P. apium,* and *L. kunkeei* decrease in relative abundance approaching the rectum, while *L. firm5, L. firm4, Bifidobacterium* and Alpha 2.1 increase (Table S3). *S. alvi* and *G. apicola* occur sporadically at low (< 1%) relative abundance throughout all queen niches.

### MANOVA of queen microbiota by chronological age and source

The two-way MANOVA performed for each of the four queen niches revealed significant variation due to chronological age, source and interaction (Table 1, Table S4). In the mouth, *P. apium* and *L. firm5* increased with age, while Alpha 2.1 and *Delftia* were more abundant in young queens (Fig. 2). The midgut and ileum aged similarly; in both niches, *Bifidobacterium* and *L. kunkeei* were more abundant in old queens while Alpha 2.1, *Delftia* and “OTHER” all decreased with age (Fig. 1). Most abundant in the ileum, *S. alvi* bloomed in 4 of 63 individuals and increased with age, while *P. apium* and *G. apicola* decreased (Table S3). In the rectum, where *L. firm5* represents the majority of total gut bacteria, *Bifidobacterium* abundance increased with age while *L. firm5* and both core Acetobacteraceae (*P. apium* and Alpha 2.1) decreased (Fig. 2). Wilcoxon rank sum tests revealed significant differences by chronological age, many of which agree with age-specific differences detected in the two-way MANOVA (Table 1, Table S4).

**Table 1.** Results examining bacterial abundance by age, niche and carbonyl accumulation.

| Category, species or OTU<sup>A</sup> |  | Abundant<br>or Rare<sup>B</sup> | Percent<br>change<br>w/age<sup>C</sup> | Wilcoxon<br>rank sum<br>test<sup>D</sup> | MANOVA<sup>E</sup> |  |
| --- | --- | --- | --- | --- | --- | --- |
| <u>Mouth</u> |  |  |  |  | <u>F value</u> | <u>Pr>F</u> |
| *Lactobacillus Firm5* |  | A | + 59 | 0.03 | - | ns |
| * *Lactobacillus Firm4* |  | R | + 195 | 0.03 | - | ns |
| *Parasaccharibacter apium* ** |  | A | + 341 | 0.02 | - | ns |
| Acetobacteraceae Alpha2.1 |  | A | - 97 | ns | 7.8 | 0.007 |
| *Snodgrassella alvi* |  | R | + 121 | 0.01 | 5.6 | 0.02 |
| *Delftia* (Burkholdariales) |  | R | - 53 | 0.0001 | 12.4 | 0.0008 |
| Diversity ($\Sigma$ OTUs 10-500) | | R | - 4 | ns | 4.5 | 0.04 |
| <u>Midgut</u> |  |  |  |  |  |  |
| *Bifidobacterium* ** |  | R | + 242 | 0.01 | 14.1 | 0.0004 |
| *Lactobacillus kunkeei* ** |  | A | + 336 | 0.01 | 5.2 | 0.03 |
| Acetobacteraceae Alpha2.1 |  | R | - 79 | 0.005 | - | ns |
| * *Delftia* (Burkholdariales)** |  | R | - 93 | <0.0001 | 26.2 | <0.0001 |
| *Diversity ($\Sigma$ OTUs 10-500) | | R | - 69 | 0.002 | 10.1 | 0.002 |
| <u>Ileum</u> |  |  |  |  |  |  |
| *Bifidobacterium* ** |  | A | + 164 | ns | 5.9 | 0.02 |
| *Lactobacillus Firm5* ** |  | A | + 9 | ns | - | ns |
| * *Lactobacillus kunkeei* |  | A | + 248 | ns | 6.4 | 0.01 |
| Acetobacteraceae Alpha2.1 |  | A | - 90 | 0.0006 | 7.1 | 0.01 |
| * *Parasaccharibacter apium* |  | A | - 50 | 0.02 | - | ns |
| * *Snodgrassella alvi* |  | A | + 279 | ns | 11.42 | 0.001 |
| * *Gilliamella apicola* ** |  | R | - 69 | 0.02 | - | ns |
| *Delftia* (Burkholdariales)** |  | R | - 94 | <0.0001 | 36.2 | <0.0001 |
| *Diversity ($\Sigma$ OTUs 10-500) | | R | - 71 | <0.0001 | 6.7 | 0.01 |
| <u>Rectum</u> |  |  |  |  |  |  |
| *Bifidobacterium* ** |  | A | + 212 | 0.001 | 15.5 | 0.0002 |
| *Lactobacillus Firm5* |  | A | - 33 | 0.03 | 9.3 | 0.004 |
| Acetobacteraceae Alpha2.1 |  | A | - 76 | 0.009 | - | ns |
| * *Parasaccharibacter apium* ** |  | A | - 78 | 0.04 | 11.7 | 0.001 |
| * *Snodgrassella alvi* |  | R | + 984 | 0.006 | 11.4 | 0.001 |
| * *Delftia* (Burkholdariales)** |  | R | - 92 | <0.0001 | 35.7 | <0.0001 |
| Diversity ($\Sigma$ OTUs 10-500) | | R | + 3 | ns | 6.3 | 0.02 |

<sup>A</sup> Dependent variables are absolute abundance of OTUs 1-9 corrected for community size (qPCR) and species 16S copy number.
Remaining OTU reads summed ( $\Sigma$ OTUs 10-500), corrected for community size and mean 16S copy number (4.2).
<sup>B</sup> Rare = < 1% mean bacterial cell number by niche (Table S2).
<sup>C</sup> Average percent change in bacterial cell number with age. We note that cell number loss cannot exceed 100%.
<sup>D</sup> Comparing species-specific bacterial cell number by chronological age only (Table S4).
<sup>E</sup> Independent variables are queen age and background (df = 3, 59). Reports only F- values for chronological age effects
Examining the top 9 most abundant OTUs and non-core diversity abundance = ( $\Sigma$ OTUs 10-500).
\* Significant interaction effect of age and background detected by the 2-way MANOVA (Table S8).
\*\* Significant MANCOVA result and Pearson correlation of bacterial abundance and carbonyl accumulation (Tables S9).

**Figure 1.**
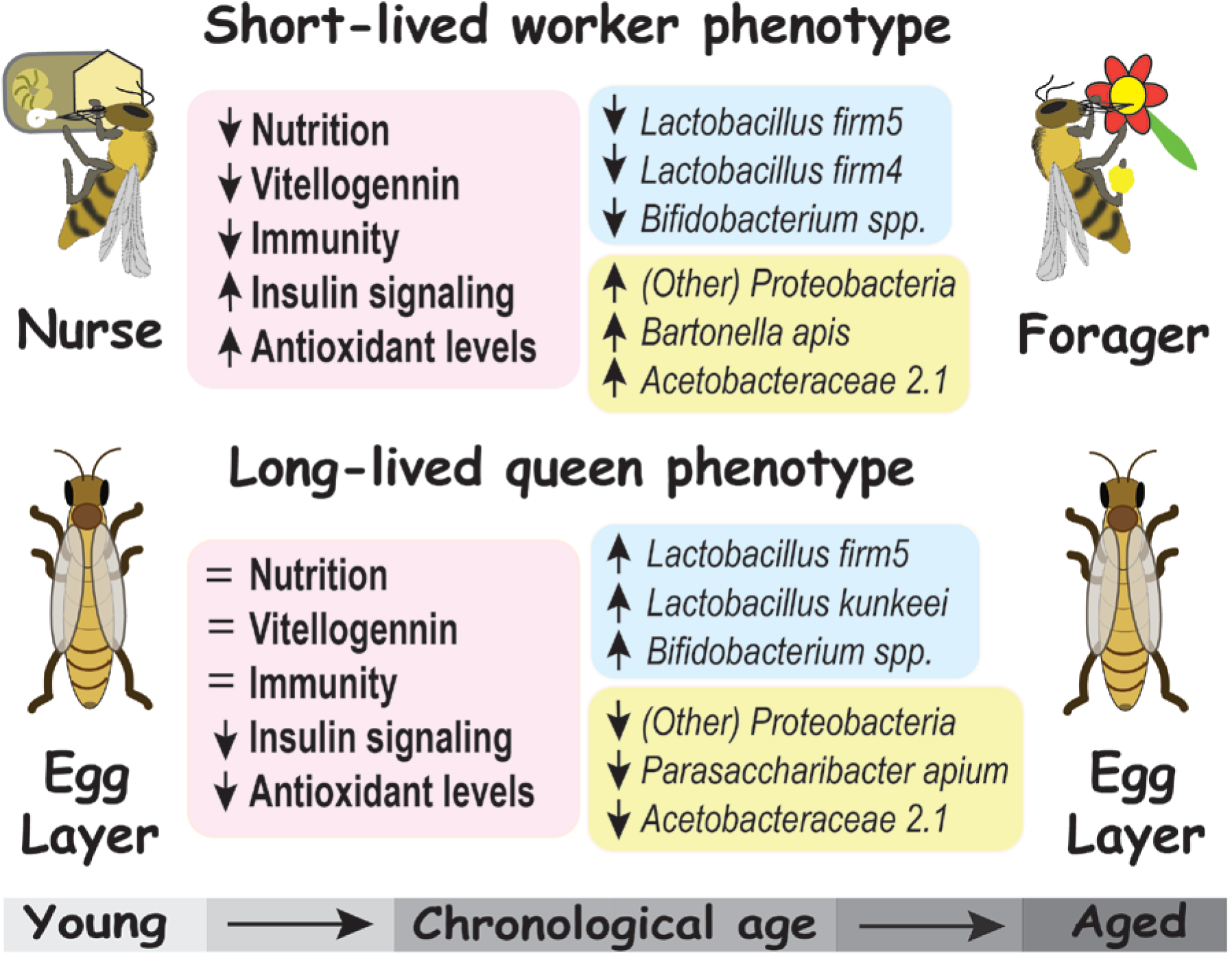
Age associated bacterial succession of extreme longevity phenotypes. Honey bee host differences (pink panels) reflect aging physiology. In the context of life history theory workers are literally the “disposable soma”, while queens represent reproduction [10]. Vertical arrows indicate the direction of change with increasing age. Firmicutes are listed in the blue panels, and Proteobacteria in the yellow panels. All listed bacterial groups differ significantly in ratio abundance. The microbiota of the short-lived worker phenotype represents a metaanalysis of *Apis mellifera* gut libraries from Kwong *et al.* [19]. Queens were analyzed in the present study (see results; Tables 1, S4).

**Figure 2.**
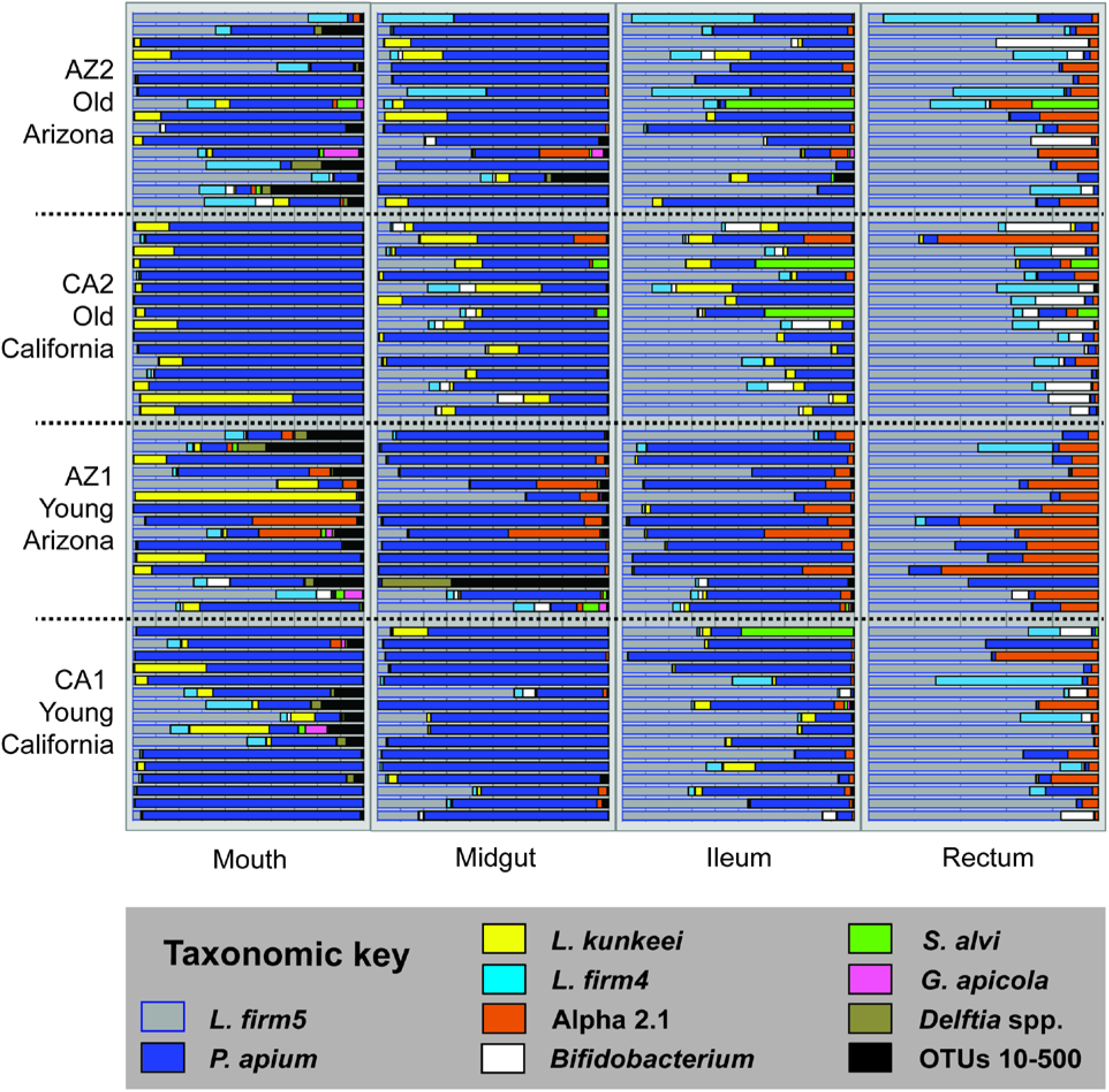
The honey bee queen microbiota by tissue. Color coded bars represents relative abundance corrected by species-specific 16S rRNA gene copy number. (See Table S3 for absolute abundance). The 4×4 panel displays the top 9 most abundant OTUs by niche, age and source. Black represents “diversity abundance”, the summation of OTUs 10-500. Old queens in the upper two rows are 16-18 months of age and young queens in the bottom two rows are aged 4.5-5.7 months (Fig. 4).

### MANOVA of worker microbiota by age

From data published in Kwong et al. (2017) we investigated age according to the behavioral role of the worker. The one-way MANOVA of microbiota by worker age (task) revealed major proportional shifts among core gut bacteria (Fig. 1). All three core Firmicutes (*Bifidobacterium, L. firm5* and *L. firm4*) decreased significantly with age while “OTHER” bacteria, Acetobacteraceae Alpha 2.1, and *Bartonella apis* increased significantly (Table S5). Of note, *S. alvi* decreased in relative abundance with age but was borderline insignificant (p = 0.06). With reference to these results, and other worker data sets in the literature we define four worker-specific gut species, all Proteobacteria, two queen-specific species, and four species shared by longevity phenotypes (Fig. 3). In general, the species shared by longevity phenotypes are particular to the rectum while the ileum species show fidelity by longevity phenotype.

**Figure 3.**
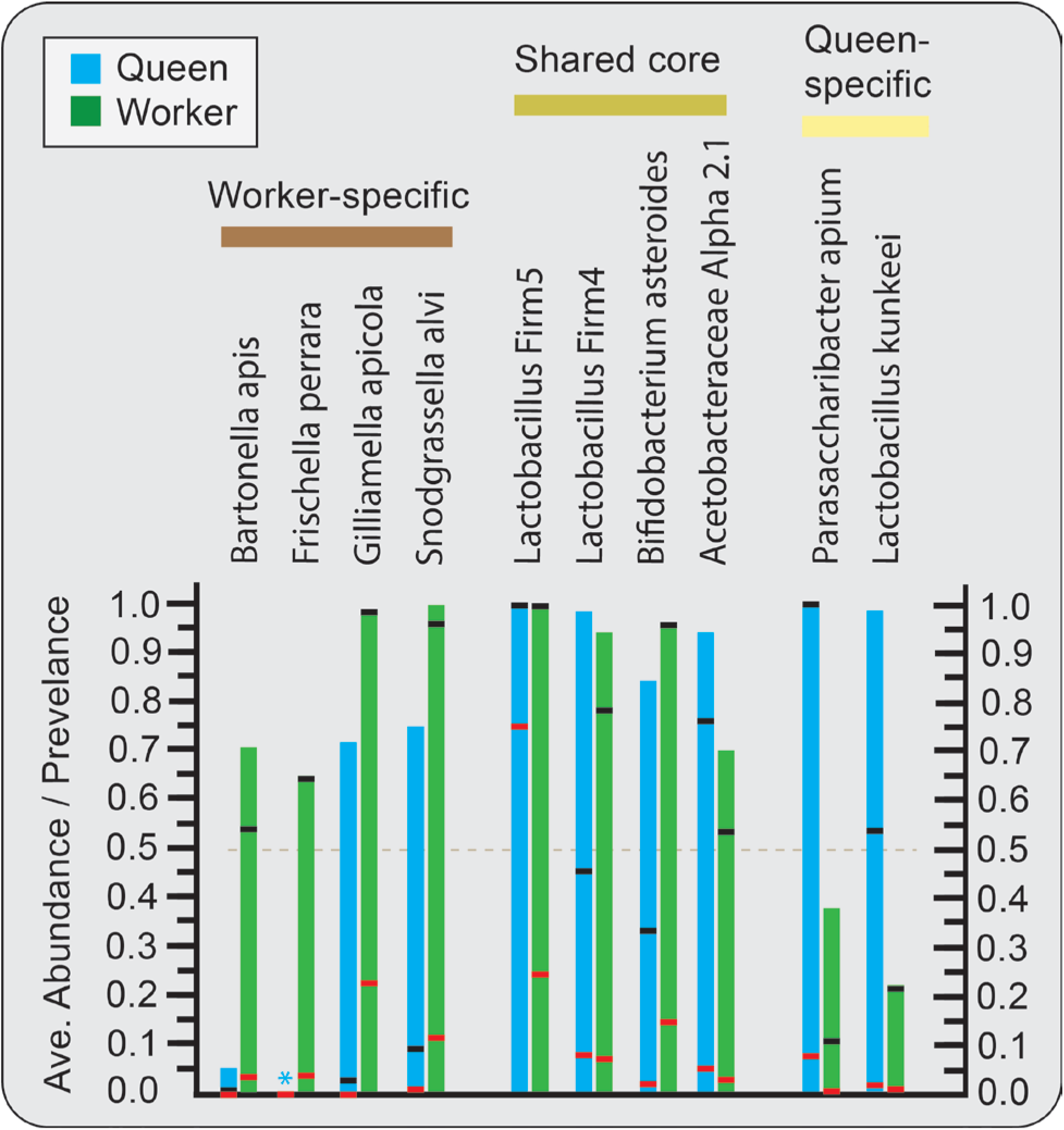
Abundance and prevalence of gut bacteria in queens (n = 63) and workers (n = 83). Workers are whole gut samples from Kwong *et al.* [19]. Queen data was normalized by tissue-specific community size to reflect relative abundance values expected from sampling whole guts. The red bars represent average abundance, black bars are prevalence defined at ≥ 0.5% relative abundance, and the bar apex is prevalence defined as 2 or more reads per gut library. We did not detect *F. perrara\** in any of the four sampled queen alimentary tract niches.

### Molecular age and the queen microbiota

We measured carbonyl accumulation in the queen fat body as a proxy for queen molecular age. While some variation in carbonyl accumulation is due to genetics and background, difficult to excrete waste products accumulate in a clock-like fashion with age. We found that chronological age was strongly associated with carbonyl content in the fat body of the queen (Fig. 4). Carbonyl accumulation differed by both age and source (Table S6). Examining all pairwise combinations, only first year queens (CA1 and AZ1) did not differ in average carbonyl accumulation. In both sets of young and old queens, chronological age did not agree with molecular age. In both age classes, queens from the Imperial Valley of California (source CA) were chronologically younger, but biologically older with greater carbonyl accumulation (Fig. 4).

**Figure 4.**
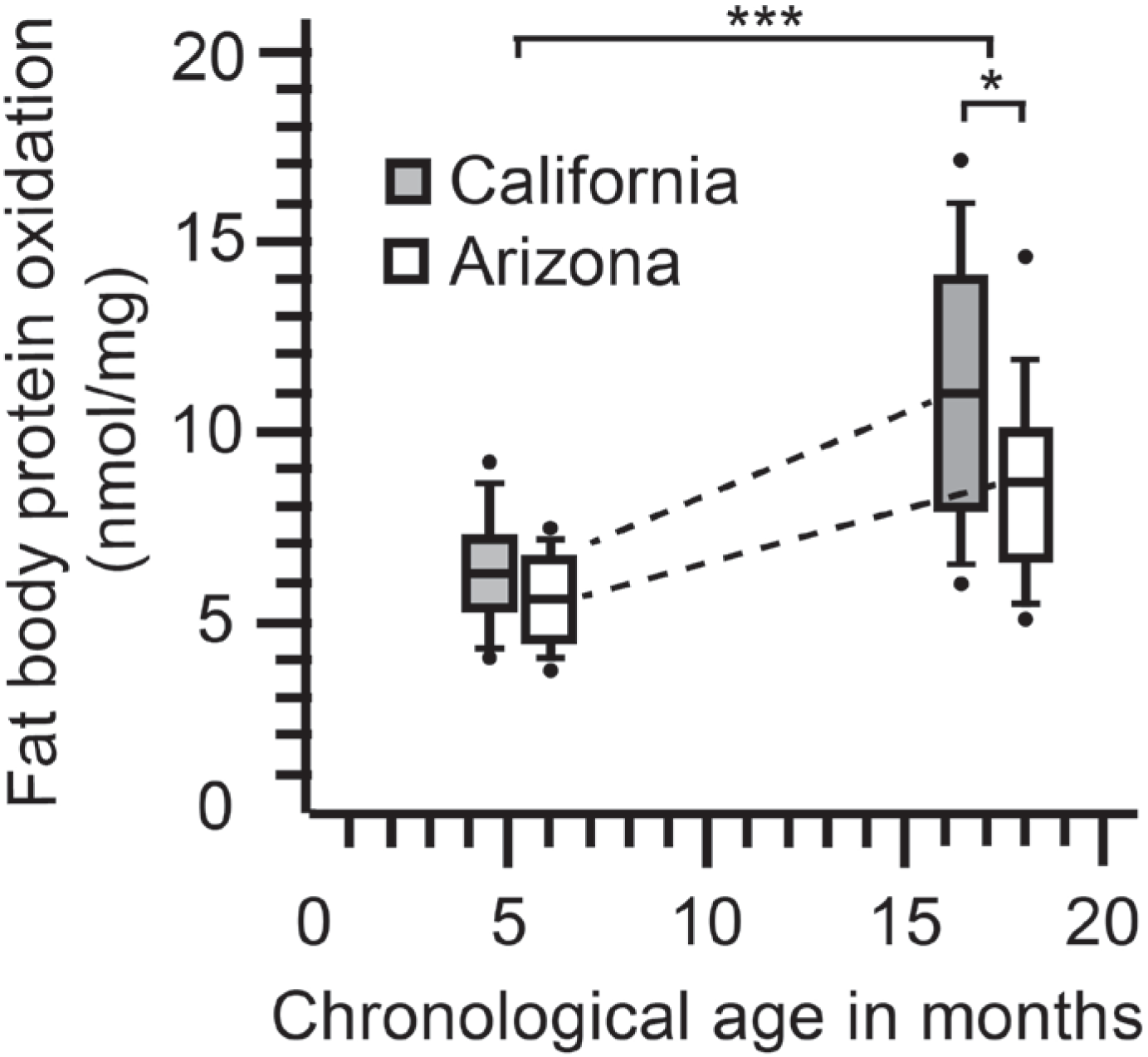
Carbonyl accumulation (protein oxidation) in queen fat body differs by chronological age (F_3, 59_ = 48.3; P < 0.0001***), and source: (t = 2.2; df = 30, P = 0.03*).

To further explore the relationship of carbonyl accumulation with queen microbiota, we ran a set of related analyses that partition variation by different strategies. Based on Bray-Curtis similarities, DistLM revealed a significant association between microbiota composition and carbonyl accumulation in each of the four tested communities (Mouth; Pseudo-F_61_ = 4.1: P = 0.01, Midgut; Pseudo-F_61_ = 3.7: P = 0.006, Ileum; Pseudo-F_61_ = 4.1: P = 0.004, Rectum; Pseudo-F_61_ = 3.9: P = 0.005). Although all four communities were significantly associated with carbonyl accumulation, little variation was explained by the collective community (mean R-sq = 0.06) due to opposing species variation within communities. The separation of background (source), chronological age and carbonyl accumulation via MANCOVA analyses detailed species-specific changes in the microbiota (Table S7). Pearson’s correlations examining species-specific CLR log transformed OTU abundance and log transformed carbonyl values agree with the main MANCOVA results examining source as the dependent variable with carbonyl accumulation as the covariate without reference to chronological time (Table S8). Most notably, throughout the gut *Bifidobacterium* is correlated significantly with the accumulation of carbonyl in abdominal fat body tissue (Fig. 5). Although at similar abundance in chronologically old and young queens, *L. firm5* abundance was also correlated strongly and positively with carbonyl accumulation. Although rare throughout the queen gut, an undescribed Burkholdariales; *Delftia* showed the strongest negative relationship with carbonyl content, decreasing dramatically with age, and varying by source (Table 1).

**Figure 5.**
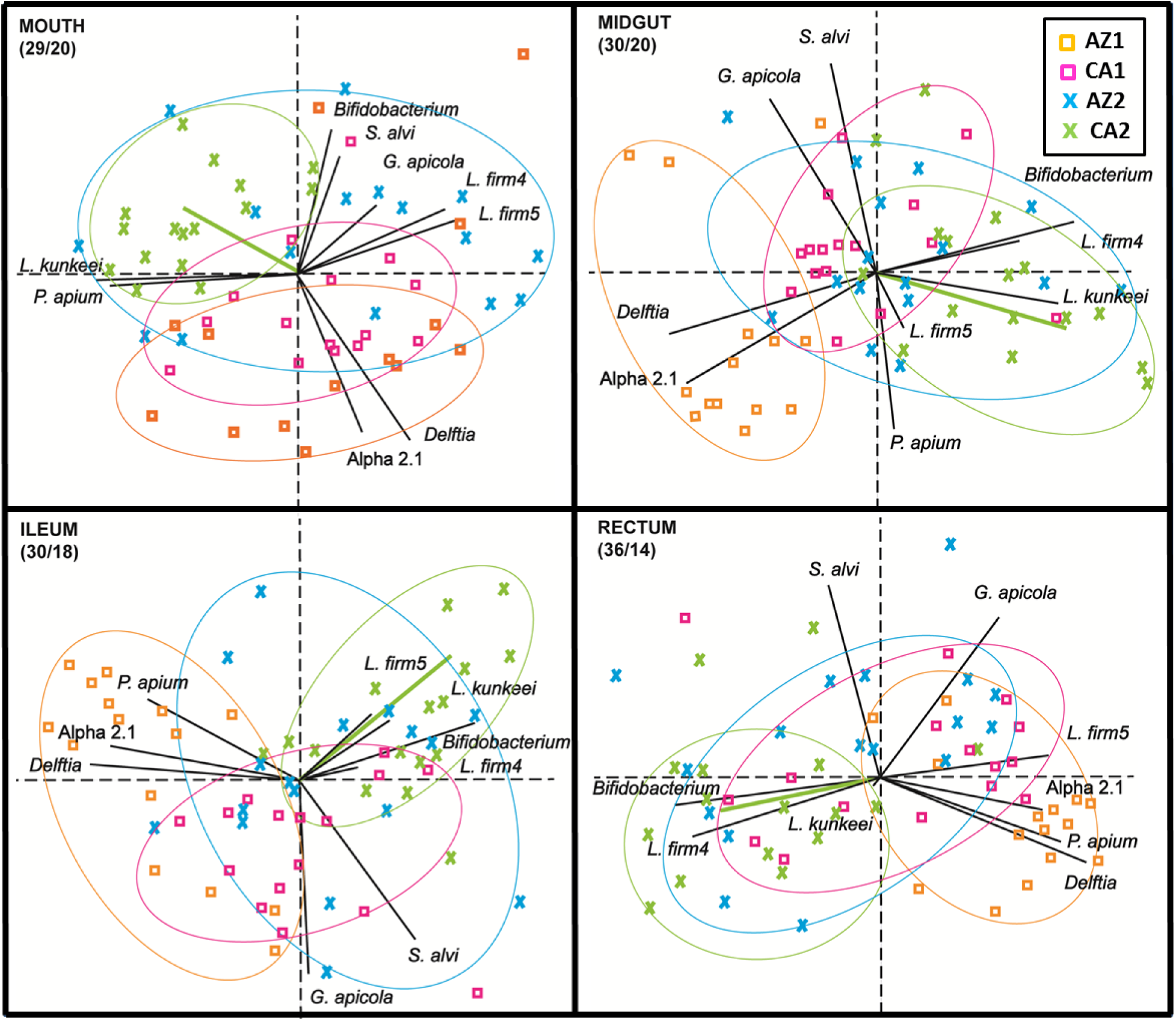
Principle components analysis by niche based on the top 9 most abundant OTUs and carbonyl accumulation. The colored symbols illustrate differences among the chronological sample cohorts; Pink and orange are young, blue and green are old. The green vector illustrates carbonyl accumulation relative to community structure, shows strong affinity with increased Firmicutes in the gut and is largely allied with the biologically oldest queen cohort (green symbols). Orange symbols are biologically youngest and consistently allied with *P. apium* in the hindgut, and Acetobacteraceae Alpha 2.1 and *Delftia* throughout the system. Biplot constructed with bacterial cell abundance data, transformed to centered log ratios (CLR) that represent the change in taxon abundance (covariance) relative to all other taxa in the data set. Thus the species vectors are proportional to the standard deviation of the ratio of each taxon to all other taxa. In general, clustered groups of points contain similar groupings of taxa with similar ratio abundances, and longer OTU vectors result from greater variation in CLR scores. The parentheses below each niche label contain the percent variation explained by the first and second principle component respectively (Table S9).

To better visualize variation associated with biological age in the queen microbiota, we performed PCA analysis using centered log ratios from the top 9 OTUs and associated carbonyl values from the fat body of each queen (Fig. 5, Table S9). Across each niche the first two principle components explained approximately 50% of the variation in log ratio abundance scores. Because the queen microbiota has shallow, deep and noisy structure, the third and fourth principle components for each niche explained an average of 14% and 8% respectively (Table S8). Although only 50% of the variation is presented in the two dimensional PCAs, a strong and consistent separation of two queen cohorts is realized in every niche; young Arizona (AZ1) and old California (CA2). In each niche, the carbonyl vector indicates CA2 as the oldest, and AZ1 as the youngest cohort, consistent with determinations of molecular (biological) age.

We examined microbiota correlations using SparCC, an approach that incorporates the structure of the data matrix to identify potential species interactions and generates null expectations based on permutation of OTU columns in the transformed data matrix. Based on SparCC, the mouth and midgut reveal a number of significant positive relationships between core bacteria within niche (Table S10). We note that SparCC results are unreliable when OTU sparsity exceeds 70% zero values but robust to communities with a low effective number of species (see Table 1). The ileum reveals a marked decrease in positive relationships, and the first occurrence of significant negative relationships. As the relevant dynamic, the two major Acetobacteraceae (*Alpha 2.1* and *P. apium*) associate positively in the ileum, but both associate negatively with *L. Firm5 and Bifidobacterium*. The strongest negative correlation occurs between *L. firm5* and *P. apium,* the two most abundant ileum species (Table S10). With more detailed investigation, age-specific Pearson’s correlations on log transformed absolute abundance shows that as queens age, the relationship of *L. firm5 / P. apium* cell number shifts from mildly negative (Pearson’s r = −0.27, p < 0.07) to strongly positive (Pearson’s r = 0.49, p < 0.005), concurrent with the loss of Acetobacteraceae (Alpha 2.1 and *P. apium*).

## Discussion

We show that host phenotypes with extreme longevity differences support gut microbiotas that age differently (Fig. 1). Because long and short-lived phenotypes are produced from the same genotype, microbiota establishment and age-associated changes likely reflect host gene expression and environmental exposure, primarily diet. Long-lived (queen) phenotypes are fed royal jelly throughout their lives to replenish internal levels of vitellogenin. In their youth, short lived (worker) phenotypes consume pollen to produce a discrete pulse of vitellogenin that fuels royal jelly synthesis in their head glands. In old age, workers forage for pollen and nectar consuming honey to support flight metabolism. This fundamental difference in diet and task reflects a suite of age-associated host gene expression, highlighted by differences in immunity, insulin signaling and antioxidant levels [9,14,16,25,52,53]. These core changes in host physiology are consistent with the distinct microbiota compositions and age-based succession of honey bee longevity phenotypes (Fig. 1). In general, aging worker guts show decreased Firmicutes and increased Proteobacteria adding to the list of insect and mammal systems where this pattern has been documented. In stark contrast, the gut microbiota of aging queens is depleted of core and other Proteobacteria, and accumulates core Firmicutes typically considered probiotic like *Lactobacillus* and *Bifidobacterium*.

### Longevity phenotypes differ in core membership

Microbiotas of long and short-lived phenotypes differ markedly in core bacterial membership sharing four of ten species, with six species showing strong phenotype-specificity (Fig. 3). The four Proteobacteria associated with worker phenotypes show distinct patterns of rarity in queen guts. In an evolutionary context, the two most recent additions to the worker gut microbiota are *B. apis* and *F. perrara* [19]. Perhaps a result of this novelty, these bacteria exhibit a relatively narrow niche breadth. *F. perrara* is specific to the worker pylorus and results in host melanization response, while *B. apis* appears in the hindguts of older foragers [28,54–56]. In queens, *B. apis* was extremely rare and *F. perrara* not detected, not even on the mouth, suggesting that these particular Proteobacteria are not tolerated by the queen, excluded via some mechanism, or result in host death. In contrast, worker-specific *G. apicola* and *S. alvi* are tolerated at low levels in queen guts, and *S. alvi* showed sporadic abundance in 4 of 63 seemingly healthy queen ileums (Fig. 2). In workers, this species pair is omnipresent, accounts for 20-60% of the ileum microbiota and represents a core syntrophic relationship critical to gut oxygen balance [25,26,57,58]. Although rare in queens, this species pair is highly correlated throughout all sampled queen niches occurring with <1% average abundance, but 70% prevalence (Fig. 3). Queen-specific bacterial species are Acetobacteraceae *Parasaccharibacter apium* and *Lactobacillus kunkeei,* both showing strong fidelity for queen mouth, midgut and ileum. These two species occur with sporadic abundance in worker ileums under conditions of putative dysbiosis and oxidative stress [12,23,29,34].

The four species shared by queens and workers differ in abundance and prevalence showing strong niche fidelity (Fig. 3). Of these four, only *Lactobacillus firm5* is core to both the ileum and rectum of both longevity phenotypes. Considering whole guts independent of age, queens and workers average 75% and 24% relative abundance of *L. firm5* respectively. Three of the shared core groups (*Lactobacillus firm5, L. firm4*, and *Bifidobacterium*) populate 100% of workers by 3-days of age [22]. Of these three, *Bifidobacterium* is significantly more abundant and prevalent in workers than queens (particularly young workers), perhaps associated with pollen consumption [24,59]. However, *Bifidobacterium* increases significantly in the hindguts of aging queens independent of pollen consumption. The fourth shared bacterium, Acetobacteraceae Alpha 2.1, is abundant in young queens but not typically detected in young workers. It decreases with queen age but becomes prevalent and abundant in older workers [19,31,60–64].

### The queen microbiota improves with age

Our results are consistent with the body of work detailing molecular aging and oxidative stress in queens, workers and social insects in general [8–10,65]. Results from the queen carbonyl assay demonstrate that queens accrue oxidative damage with age (Fig. 4), and that chronological age can differ significantly from biological age possibly due to environmental differences including climate, nutrition, toxins and other landscape variables. Despite similar signs of biological aging in both queens and workers, the gut microbiota of older queens seems to reflect a refined structure with greater efficiency. It’s unlikely that queens ever develop a senescence physiology and associated microbiota as seen in workers. Under natural conditions, queens accrue molecular damage associated with aging, but are not allowed to grow “old” because fecundity is critical to colony survival, and workers routinely replace substandard queens [66]. With increased oxidative damage, gram positive bacteria decrease in workers [19] but increase in queens (Fig 1). Of note, core *Lactobacillus* and *Bifidobacterium* in queens show greater correspondence with biological than chronological age suggesting that these species may track or signal host physiology (Table S7). Consistent with decreased antioxidant expression and less ROS generation in queens [9,65] the bacteria that increase with queen age do not rely on oxygen, but generate continuous fermentative metabolism in the queen hindgut (Fig. 2). In turn, these fermentation products (i.e. butyrate) are considered fundamental to host physiology and homeostasis [25,59].

Results from conventionalized bee experiments suggest that butyrate produced by the honey bee hindgut microbiota plays a key role in host metabolism [25,59]. In human colons, positive butyrogenic effects are considered a result of cross feeding by butyrate-producing Firmicutes and *Bifidobacterium* [67]. Feeding worker honey bees relevant amounts of sodium butyrate results in gene expression considered beneficial to general health, broadly affecting immunity and detoxification [68]. We found that bacterial communities implicated in butyrate production were diminished in aging workers but seemingly enhanced in aging queens. Better explained by biological than chronological age (Fig. 5), *Bifidobacterium* increases significantly with age in the queen midgut, ileum and rectum (Table S7). Moreover, *L. kunkeei* increases in the midgut and ileum, while community changes in the ileum favor the persistence of *L. firm5* (Table 1), and suggest a more efficient relationship emerges with queen age. *Lactobacillus firm5* is the most plentiful bacteria in the queen hindgut, and combined with increased *Bifidobacterium*, may add to the butyrogenic effect in queens concurrent with increased biological age. *Bifidobacterium* itself was recently identified as a major bacterium associated with host-derived signaling molecules in worker honey bees [59]. Interestingly, *Bifidobacterium* abundance in both queens and workers is often low and/or highly variable so may be affected by diet or strain variability [24].

We compared the gut microbiota of young in-hive bees to older foragers within and among studies. Foraging is the last functional role workers serve before death. But as a group, both in-hive bees and foragers can range greatly in chronological age and environmental exposure [69]. Also, comparing across next generation sequencing studies can be misleading due to differences in methodology like primer choice or analysis pipeline [70]. Despite these and other sources of potential error, we found that the worker gut microbiota ages in a highly predictable fashion, becoming depleted of core hindgut firmicutes including *Bifidobacterium* (Fig. 1). Of seven available forager studies, three used the same methods to sequence both foragers and nurses [19,31,63], and we used these studies as a point of reference for examining worker aging. We analyzed the largest and most variable of these three data sets [19] defining six significant differences in microbiota between young and old workers (Fig. 1). The collective results from six of seven studies are largely in agreement and suggest that age-associated shifts in worker microbiota are strongly predictable at the level of species despite study particulars [19,31,60–64]. The changes we report in figure 1 [19] represent a functional change from a fermentative to proteolytic hindgut environment, involving significant shifts in core bacterial structure. Alpha 2.1 increases in all 7 studies*, B. apis* and “Other” bacteria in 6 of 7. One to three major core hindgut Firmicutes are depleted significantly in 6 of 7 studies, while studies were more variable concerning shifts of *S. alvi* and *G. apicola*, the species pair that dominates the worker ileum.

### Early gut succession

Similar to workers [37], the rectum of the mature queen contains 84% of the total bacteria found in the queen gut (Fig. 2). On average, a whole gut sample from a mature laying queen would be highly biased toward rectum species, dominated by *Lactobacillus firm5* (Fig. 3). In contrast, whole gut samples of queens during the mating process show a dominant Acetobacteraceae (*P. apium* and Alpha 2.1) profile [20]. This finding is consistent with our detection of significantly more *P. apium* and Alpha 2.1 in the guts of younger queens (Table 1), and suggests that the bacterial succession leading to a *L. firm5* dominant hindgut in queens may require many weeks, perhaps months. Given that worker gut succession occurs throughout the life of the worker [12,22], we speculate that the early queen microbiota [20] represents a pioneer community that primes the gut environment or host immune system, and/or potentially aids disease prevention during the days-long mating process that involves queen flight metabolism and mating with >20 males. A successfully mated queen is fed massive amounts of royal jelly as she begins to lay eggs. The decrease and stabilization of cell replacement rate in early queen midguts [71] suggests a more stable gut environment emerges around 40 days of age, perhaps influencing bacterial succession.

### Queen niche breadth

In queens, the occurrence patterns and numerical dominance of *P. apium* in the mouth and midgut, and *L. firm5* in the ileum and rectum suggests that the extended gut structure is important for host function (Fig. 3). The taxonomic shift at the pylorus demarcates a steep physiological gradient in the adult bee gut. Recently characterized in workers, this change occurs just upstream of the ileum where Malpighian tubules feed host waste products back into the gut, and microoxygenic and pH gradients affect bacterial establishment and persistence [25,29,54]. Host excretions provide a different niche for bacterial co-evolution including an influx of nitrogenous waste compounds, a decrease in oxygen availability and lower pH [25]. While the effect of pollen consumption on host signaling has been investigated in workers [24,25], the effect of the queen’s diet (royal jelly) on host signaling remains unknown. The reliable and predigested nature of the queen diet may generate very different collection of waste products, supporting hindgut bacterial strains distinct from those found in workers.

It is mostly unknown why queens can resist many worker diseases and vice-versa. Early queen death has become more common [66,72], and defining disease states in queens will rely in part on the structure and function of native gut bacteria [12]. Although rare throughout the gut, the occurrence pattern of *Delftia* (Burkholdariales) suggests it is detrimental. Not detected in workers, *Delftia* is negatively correlated with *L. firm5* and *Bifidobacterium* in the queen hindgut, shows the greatest negative correlation with carbonyl accumulation, and decreases significantly with biological age (Table1). Congruently, *Delftia* is negatively correlated with putatively beneficial bacteria on the queen mouth and midgut (Tables S9 and S10). These two niches are dominated by distinct sequovars of *P. apium*, a bacterium co-evolved to thrive on royal jelly [7]. Over 95% of the mouth/midgut bacteria classify as *P. apium and L. kunkeei,* both associated with decreased abundance of honey bee-specific disease caused by bacteria and microsporidia [73,74]. One primary function of microbes in the queen mouth and midgut may be the exclusion of opportunistic and disease causing microbes. Mouth communities not dominated by *P. apium* are much smaller in size and contain significantly more *Delftia*, OTU diversity and “other” bacteria (Tables S3) suggesting that *P. apium* dominance in the queen mouth and midgut limits the occurrence of detrimental bacteria in the hindgut. Older queens have significantly more *P. apium* on their mouths and *L. kunkeei* in their midguts that may accrue with age and/or improve queen hygiene promoting queen survival (Fig. 2). Pollen exposure and consumption may render workers more vulnerable than queens to frequent pathogen invasion. The queen and her constant diet of royal jelly may discourage novel microbial acquisition and provide a strong selective environment for the evolution of niche specialists. The constant diet of royal jelly likely represents a form of purifying selection, perhaps even an arms race at the front end of the queen, producing fierce competition among *P. apium* strains for this constant and complete nutrient source.

### Evolution of “queen-specific” gut bacteria

*P. apium, L. kunkeei*, and close ancestors occur throughout solitary and social bees and may even predate the evolution of corbiculate bees [19,75–79]. Both *P. apium* and *L. kunkeei* grow at extreme sugar concentrations and royal jelly enhances the invitro growth of some strains [7,33]. The evolution of host behavior to mechanically concentrate nectar sugars via evaporation was likely a key innovation producing strong selection for these two osmotolerant symbionts. Bacteria adapted to survive in concentrated nectar of solitary bee provisions were well positioned to develop greater fidelity with the host gut. The mature worker ileum is dominated by core bacteria *S. alvi* and *G. apicola* that co-occur in a biofilm with lesser amounts of *Lactobacillus* firm5 [12]. In contrast, the mature queen ileum is dominated by *Lactobacillus* firm5 that co-occurs with lesser amounts of core gut bacteria *P. apium* and *L. kunkeei* (Fig 1). Reciprocally, worker ileum bacteria *S. alvi* and *G. apicola* are found at similarly low levels in the queen ileum and show sporadic abundance in the queen. These symmetrical occurrence patterns suggest antagonistic co-evolution of caste-specific gut bacteria, a hypothesis consistent with host age phenotype and development-specific pathogen strategies.

Over 16 *L. kunkeei* genomes have been compared, revealing core functionality and a large variety of accessory protein clusters that characterize different strains [80]. Isolated from the gut of *A. mellifera,* strains MP2 and EFB6 of *L. kunkeei* were most related, and differ from other *L. kunkeei* in possessing genes implicated in gut colonization including cell adhesion, biofilm formation and horizontal transfer [35,80]. These likely represent strains that colonize the queen midgut and ileum. They may also colonize gut environments of workers and larva. That many of the *L. kunkeei* genomes lack gut-specific genes suggests they may lead more opportunistic life cycles within the hive and pollination environment. Similarly, the genome of *Parasaccharibacter apium* also reveals multiple functional traits for biofilm life in the insect gut, including survival in low oxygen environments and adhesion to host epithelium [44]. Like *S. alvi*, and many other AAB, *P. apium* can assimilate major fermentation byproducts generated by neighboring bacteria. Collectively this suggests that *P. apium* metabolism in the queen ileum may be somewhat analogous to *S. alvi* function in the worker ileum [57].

Patterns of species co-occurrence suggest selection pressure for honey bee gut bacteria to co-exist with other bacteria in a biofilm encouraging competition and co-evolution (Fig. 4). This hypothesis is supported by the complex of highly correlated bacteria on the queen mouth, and strongly affiliated species pairs occurring regardless of niche. Not strongly associated with age, niche or background, at least three pairs of co-occurring species emerge as potential syntrophic relationships throughout the queen microbiota, and may rely on co-evolved traits to ensure niche occupation. This strategy would prove more effective in the queen gut, which provides a more stable long term environment where partnerships have more generational time to evolve. Many bacterial pairs have evolved strict affiliations with one another and multiple hive niches including *P. apium / L. kunkeei, G apicola / S. alvi* and *L. firm4 / Bifidobacterium* (Figs. 3 and 4). Perhaps through their reliance on one another, core bacteria better survive within and outside their preferred niche.

## Conclusions

The honey bee is a metabolic model for the effects of aging and diet on microbiota. Sampling the honey bee microbiota with respect to chronological age, biological age and environmental exposure facilitates an informative partitioning of variation associated with longevity phenotypes. Consistent with research on aging and host oxygen dynamics, the queen microbiota shifts towards the fermentative metabolism of well-known gram positive species, while the more rapidly aging worker is progressively depleted of these same species. Given the spectrum of influence of gut microbiota on worker physiology, we suggest that the queen microbiota serves a similarly critical role in host signaling and protection. Separate evolutionary trajectories for caste-specific gut bacteria reflect overt differences in diet and longevity between workers and queens. This trajectory appears to have tracked division of labor evolution, perhaps involving key innovations like nectar concentration to produce honey, and the production of royal jelly in worker hypopharyngeal glands. Once considered bacteria associated with worker gut dysbiosis and larval nutrition, *L. kunkeei* and *P. apium* must now be understood as core gut bacteria of *Apis mellifera* queens. Our results suggest that these two species occupy a functional niche in the queen mouth, midgut and ileum. The co-occurrence and correlational abundance of multiple core species throughout the honey bee system suggest syntrophic relationships are commonplace. More generally, our study highlights the importance of controlled temporal and tissue-specific data to understand the total diversity and function of the honey bee microbiome.

## Abbreviations

16S rRNA gene: 16S subunit of the ribosomal RNA gene
ANOVA: Analysis of variance
AZ: Arizona
BCA: bicinchoninic acid
BLAST: Basic local alignment search tool
bp: Base pairs
CA: California
CLR: Centered log ratio
DistLM: distance-based linear model
DNA: Deoxyribonucleic acid
DNPH: 2,4-dinitrophenylhydrazine
FDR: false discovery rate
GLM: General linear models
MANOVA: Multivariate analysis of variance
MANCOVA: Multivariate analysis of covariance
OTU: Operational taxonomic unit
PCA: Principal Components Analysis
qPCR: quantitative polymerase chain reaction
RDP: Ribosomal Database Project
rRNA: ribosomal ribonucleic acid
SparCC: Sparse Correlations for Compositional data algorithm

## Declarations

### Ethics approval and consent to participate

Not applicable

### Consent for publication

Not applicable

### Availability of data and material

Honey bee queen datasets were deposited with the NCBI BioProject database. BioProject ID: PRJNA438524, https://www.ncbi.nlm.nih.gov/Traces/study/?acc=SRP135870

Summary list view available here: https://www.ncbi.nlm.nih.gov/sra/?term=SRP135870

Honey bee worker dataset can be accessed here: https://doi.org/10.1126/sciadv.1600513

### Authors’ contributions

K.E.A. conceived of and designed the research. V.A.R. and B.M.M. contributed new analytical tools. V.A.R., B.M.M., D.C.C., A.C.F. and P.M. performed the experiments. K.E.A. V.A.R. and P.M. analyzed the data. K.E.A. wrote the manuscript. All authors read and approved the final manuscript.

## Acknowledgements

The corresponding author thanks Belynda Starr, Ariel Calypso and Isaak El Spaghett for their valuable input. We thank the BIO5 institute at the University of Arizona for amplicon sequencing, and two anonymous reviewers for providing helpful comments on the manuscript. The USDA/ARS is an equal opportunity employer and provider.

